# Hox gene expression during the development of the phoronid *Phoronopsis harmeri*

**DOI:** 10.1101/799056

**Authors:** Ludwik Gąsiorowski, Andreas Hejnol

## Abstract

**Background:** Phoronida is a small group of marine worm-like suspension feeders, which together with brachiopods and bryozoans form the clade Lophophorata. Although their development is well studied on the morphological level, data regarding gene expression during this process are scarce and restricted to the analysis of relatively few transcription factors. Here we present a description of the expression patterns of Hox genes during the embryonic and larval development of the phoronid *Phoronopsis harmeri*.

**Results:** We identified sequences of 8 Hox genes in the transcriptome of *P. harmeri* and determined their expression pattern during embryonic and larval development using whole mount *in situ* hybridization. We found that none of the Hox genes is expressed during embryonic development. Instead their expression is initiated in the later developmental stages, when the larval body is already formed. The Hox genes are expressed in the metasomal sac, posterior mesoderm and junction between midgut and hindgut - structures that represent rudiments of the adult worm, which emerges through the process of drastic metamorphosis. Additionally, two Hox genes are expressed in the posterior telotroch, which develops in the later larval stages.

**Conclusions:** The lack of Hox gene expression during early development of *P. harmeri* indicates that the larval body develops without positional information of the Hox patterning system. Such phenomenon might be a consequence of the evolutionary intercalation of the larval form into an ancestral, direct life cycle of phoronids. Accordingly, the specific actinotrocha larva found only in Phoronida, would represent an evolutionary novelty, for which an alternative molecular mechanism of antrerior-posterior patterning was recruited. Another explanation of the observed Hox gene expression is that the actinotrocha represents a “head larva”, which is composed of the most anterior body region that is devoid of Hox gene expression. This implies that the Hox patterning system is used for the positional information of the trunk rudiments and is, therefore, delayed to the later larval stages. Future investigation on head-specific genes expression is needed to test this hypothesis.

## Background

Hox genes encode a family of transcription factors present in Bilateria and Cnidaria(1-4), which bind with their conserved homeodomain directly to regulatory regions of downstream genes and activate or suppress their expression (e.g. (5-7)). In many clades, Hox genes are differentially expressed in the early developmental stages along the anterior-posterior axis of the developing embryo, being one of the important components of molecular patterning of axial identities(4-6, 8-10). The diversity of Hox genes present in extant Bilateria originated likely by multiple duplication events, which resulted in the physical linkage of Hox genes in the genomes of many Bilateria, the so called Hox clusters (e.g. (9, 11, 12). It is possible to discriminate organized, split and disorganized Hox clusters, depending on the level of their organization(7, 12) and in certain Bilateria the Hox genes are expressed in roughly the same order as they are located in the cluster: a phenomenon referred to as collinearity(6, 9, 11). The correspondence between position of the gene in cluster and onset of its expression might have a temporal (during development) or spatial (along body axis) character and accordingly it is possible to discriminate between the temporal and spatial collinearity. It has been proposed that collinearity, especially the temporal one, is a major factor responsible for conservation (or maybe even formation) of the ordered Hox cluster in the genome(9, 11-16).

Although expression of Hox genes has been described during embryonic and larval development of many animals representing diverse evolutionary lineages(4, 16-49), there are still some clades for which information about Hox expression during development is lacking. Among them are phoronids, marine, sessile worms, which feed using a specialized filter apparatus, the so-called lophophore (*lp* in Fig. 1B). Due to the presence of lophophore, Phoronida have been traditionally united with two other clades – Ectoprocta (Bryozoa) and Brachiopoda – into the group called Lophophorata(50, 51), which recently gained support as a valid clade from several transcriptomic and phylogenomic studies(52-55). Although originally the Lophophorata were considered as deuterostomes(50, 51), molecular data showed their protostome affinity(56) and currently the lophophorates occupy a well-supported position within the clade of Spiralia(52-55, 57). Phoronids develop through a distinctive planktotrophic larval stage, called actinotrocha(58-61). After a prolonged planktonic life, the actinotrocha larva settles (Fig. 1C) and undergoes drastic metamorphosis, during which the rudiment of the body wall of the adult worm, the so called metasomal sac (ms, Fig. 1C), is everted and the rudiments of the adult internal organs descent from the larval body to the newly formed juvenile worm (Fig. 1C)(60, 61). The only exception from this pattern is *Phoronis ovalis*, which is a sister group to the remaining phoronids(62-64) and which develops through the creeping slug-like larva(60). After few days of development the active larva settles and acquires a smooth hemispherical shape(60). Unfortunately, the degree of the metamorphosis-related remodeling of internal structures in *Ph. ovalis* remains poorly examined.

**Fig. 1.**
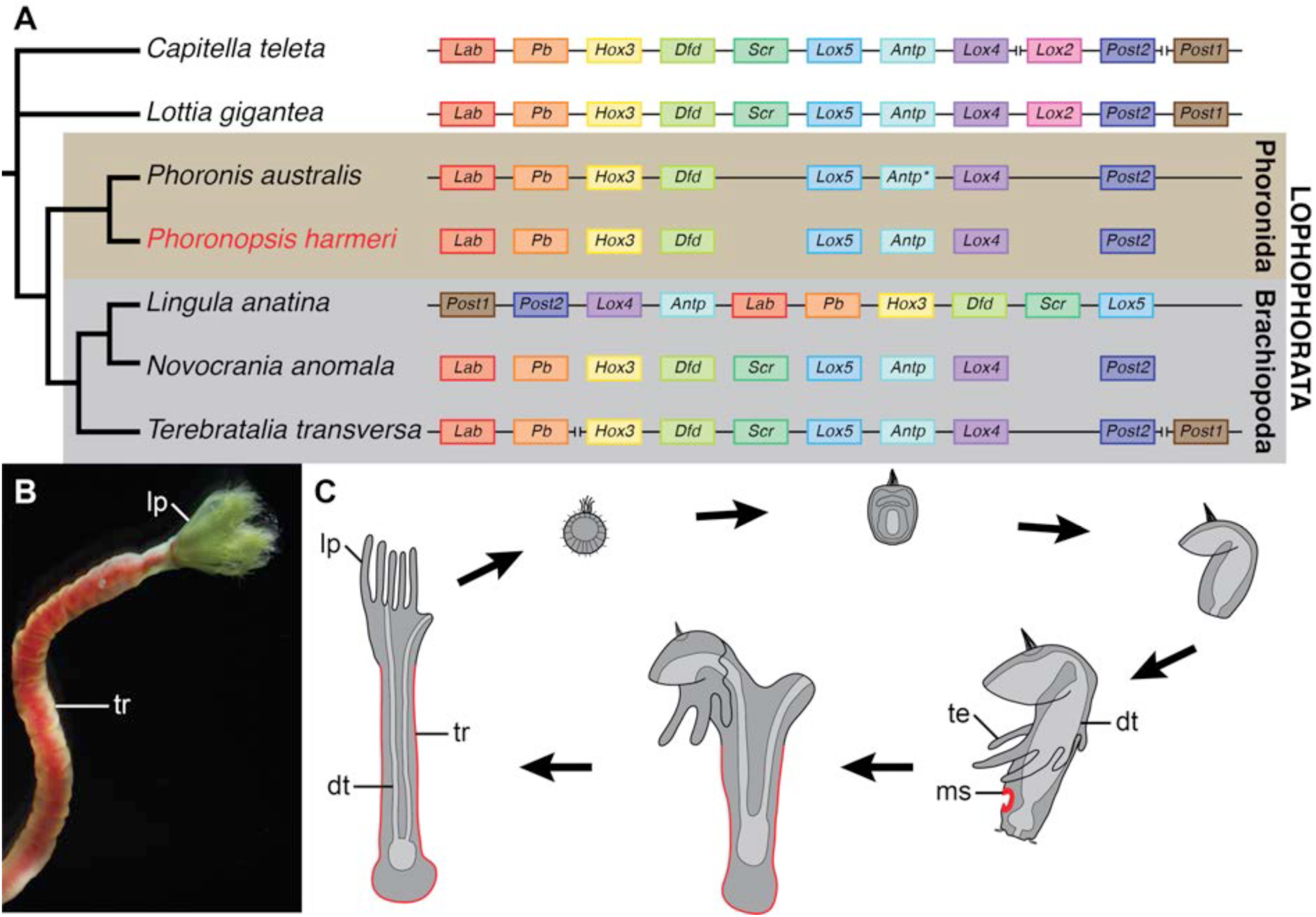
Hox gene cluster organization in various Spiralia (A), *Phoronopsis harmeri* gross morphology (B) and details of its life cycle (C). Hox cluster organization and Hox gene complement in A are taken from (16, 89, 91). Gene *antp* from *Phoronis australis* (marked with asterisk) was originally described as *lox2* (see text for discussion). Metasomal sac and adult trunk originating from it are marked in red in C. Abbreviations: *dt* digestive tract, *lp* lophophore, *ms* metasomal sac, *te* larval tentacles, *tr* trunk.

The phoronid development has been well studied on the morphological level (e.g. (58-61, 65-85)), including preliminary cell lineage, blastomere ablation and fate mapping studies(86-88). However, information about the molecular patterning is limited to the single study of 9 transcription factors (which include anterior, posterior and endomesodermal markers) during the development of *Phoronopsis harmeri*(85). Importantly, information about expression of Hox genes during development of any phoronid species is still lacking(40, 59). Recently, Lou et al. have demonstrated that in phoronid *Phoronis australis* a Hox cluster is highly organized with all of the 8 phoronid Hox genes forming a single cluster that retains the ancestral spiralian order of genes ((89), also Fig. 1A). This is in contrast to brachiopods, the putative close relatives of Phoronida, where various level of Hox cluster disorganization is shown (Fig. 1A) and temporal and spatial collinearity is missing (16, 40, 89, 90). Therefore, it remains important to examine whether phoronid Hox genes are also expressed in the spatio-temporally collinear manner during development, which would correspond with the retention of the organized Hox cluster shown in this clade.

Phoronids exhibit a biphasic life cycle with planktotrophic larvae that transform into juvenile in the catastrophic metamorphosis event (Fig. 1C; e.g. (59, 60, 73, 75, 81, 82)), the process which is much more drastic than relatively gentle metamorphosis of most Spiralia. Importantly, the A-P axis of the larva is profoundly altered during metamorphosis(60, 77, 81, 82) and results in the U-shaped organization of the internal structures of the juvenile worm (Fig. 1C). In animals with pronounced metamorphosis Hox genes might exhibit noticeable differences in the expression patterns during development of larval and adult bodies. In pilidiophoran nemerteans and indirectly developing hemichordates it has been demonstrated that Hox genes are involved in patterning of only adult bodies(37, 38), while in tunicates and sea urchins a different sets of Hox genes are expressed during larval and adult body development(21, 22, 44, 47). On the other hand, in animals with more gradual metamorphosis (e.g. cephalochordates, mollusks, annelids or brachiopods), the Hox genes seem to pattern both the larval and adult body plans in a relatively similar way(31, 39, 40, 46, 48). However, studies focusing on metamorphosis-related differences of Hox gene expression in Bilateria are still limited to a relatively few evolutionary lineages(40, 92). Therefore, the comparison of Hox gene expression between the embryonic and larval development and the development of the metasomal sac in phoronids might shed new light into the understanding of the evolution of differential genetic control of the axis patterning in animals with extreme metamorphosis.

In this study we investigated the Hox genes complement and their expression patterns during the development of the phoronid *Phoronopsis harmeri*, for which the extensive data on the morphological aspects of the development and some molecular data on the A-P axis are available(66, 72, 75-78, 80-82, 84, 85)). Our aim was to answer whether phoronid Hox genes show staggered expression along the A-P axis at any of the developmental stages as well as to examine if there are traces of temporal collinearity that could hint to the presence of a Hox cluster as described for another phoronid *Ph. australis*(89). We also wanted to investigate whether there are differences in the Hox gene expression (and possibly in the patterning of the A-P axes) between the larva and the rudiment of the forming juvenile worm and compare our findings with other species that exhibit extreme metamorphosis.

## Results

### Hox complement and gene orthology

We identified 8 Hox genes in the transcriptome of *P. harmeri* and our phylogenetic analysis allowed their assignment to particular orthology groups (Fig. 2). Those genes represent orthologues of the genes *labial* (*lab*), *proboscipedia* (*pb*), *hox3, deformed* (*dfd*), *lox5, antennapaedia* (*antp*), *lox4* and *post2* (Figs. 1A and 2). Moreover, in addition to the gene *cdx* reported by Andrikou et al.(85), we identified two other paraHox genes in the transcriptome of *P. harmeri* – *gsx* and *xlox*. Most of the Hox orthologues form distinct clades in our phylogenetic tree (Fig. 2). Sequences from the two orthologues (sex combs reduced (*scr*) and *antp*) do not form clades but rather grades of similar sequences (Fig. 2), which nevertheless allow exact orthology assessment. We found that the gene identified by Luo et al. as *lox2* in the genome of *Ph. australis*(89) and its orthologue in *P. harmeri* do not fall into the clade containing *lox2* sequences from other Spiralia, but instead they group in the grade containing *antp* sequences. Accordingly, sequence of those two phoronid genes lack most of the residues proposed as signature of *lox2* by de Rosa et al. (Additional File 1: Fig. S2; (93)).

**Fig. 2.**
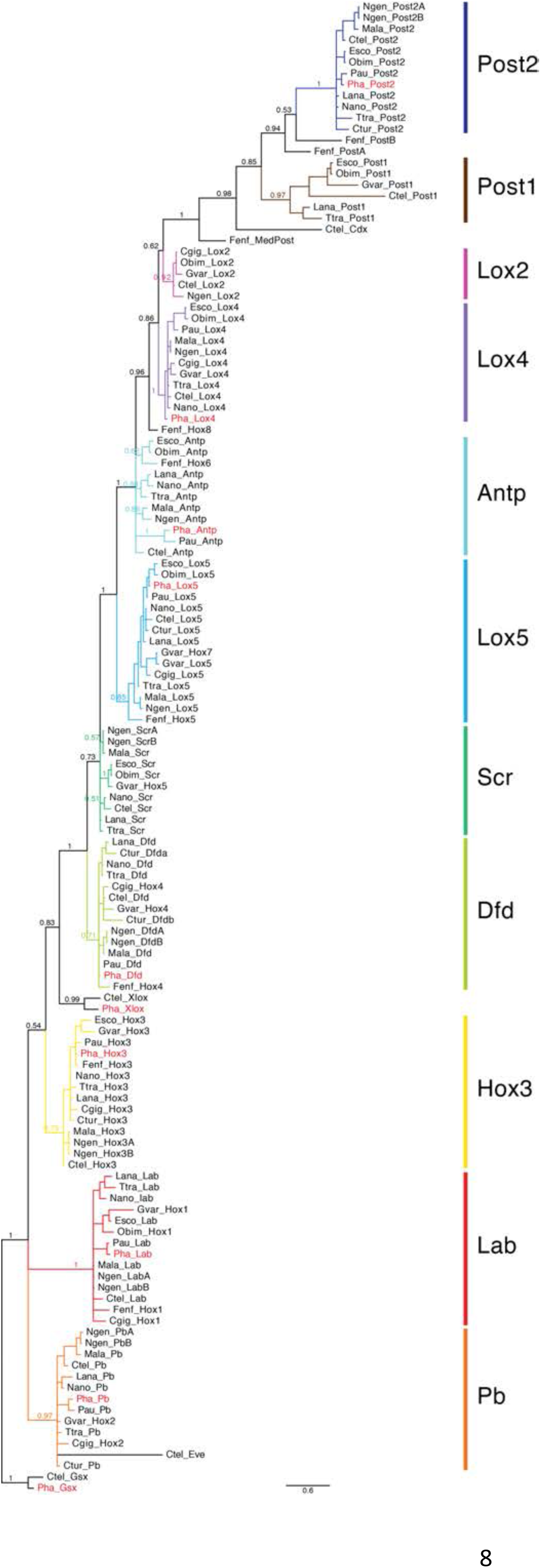
Bayesian phylogeny under JTT+I+G substitution model of the amino acid sequences of spiralian Hox genes homeodomains, including phoronid sequences. Genes from *P. harmeri* are marked in red. Posterior probability values are shown for important clades. Full species names and sequences accession number are provided in Additional file 2: Tab. S1.

### *Embryonic and larval development of* P. harmeri

Embryos and larvae of *P. harmeri* are relatively transparent and many aspects of their morphology can be easily observed with the light microscope using Nomarski interference contrast (Fig. 3). At 9°C the blastula stage is reached at about 6-8 hours post fertilization (hpf). Around 12 hpf a swimming blastula with a large blastocoel (*bc*) is formed (Fig. 3 A, A’). At 20 hpf the gastrulation process is initiated, which leads to the formation of the gastrula (Fig. 3 B, B’) that displays a distinctive blastopore (*bp*), the archenteron (*ar*) and the anterior mesoderm (*am*). Subsequently, the embryo (including the archenteron) elongates along the A-P axis and the oral hood (*oh*) develops anteriorly leading to the formation of the early larval stage, at approximately 40 hpf (Fig. 3C, C’). In the posterior part of the early larval gut the intestine (*in*) develops, which merges with the posterior ectoderm and forms an anus. Laterally to the intestine the first undifferentiated rudiments of the protonephridia are present (*pr* in Fig. 3C, C’). At 60 hpf the pre-tentacle larval stage is reached (Fig. 3D, D’), which possesses a tripartitioned digestive tract (with esophagus, *es*; stomach, *st* and intestine, *in*), an apical organ (*ao*), protonephridial rudiments (*pr*) and rudiments of the first three pairs of tentacles (*rt*). 3 days post fertilization (dpf) larvae can be already identified as early 6-tentacle actinotrochae (Fig. 3E, E’) due to the presence of three pairs of well-defined tentacles (*te*). At this stage the larval protonephridia reach their definite branching form (*pn*, Fig. 3E), the rudiments of posterior mesoderm are morphologically distinguishable (*pm*, Fig. 3E) and the posterior telotroch starts to form around anal opening (*tt*, Fig. 3E’). At 5 dpf (Fig. 3 F, F’) the telotroch is fully formed, while the posterior mesoderm forms rudiments of the posterior coelom compartment (metacoel), displacing the terminal organs of larval protonephridia to the more anterior position. The actinotrocha reach the 8-tentacle stage at 7 dpf (Fig. 3G, G’). At this stage the post-tentacular region of the body elongates and the metasomal sac, a rudiment of the body wall of the prospective adult worm, is formed (*ms*, Fig. 3G, G’). The metasomal sac at this stage appears as an ectodermal thickening located on the ventral side between tentacle bases and the telotroch. The thickening is concave and it is possible to discriminate between its opening (which will represent the anterior part of the adult trunk), and bottom (which after eversion will contribute to the posterior ampulla). The morphological details of the embryonic and larval development of *P. harmeri* are well described elsewhere(66, 72, 75-78, 80-82, 84, 85), therefore we did not examined further the embryonic and larval morphology.

**Fig. 3.**
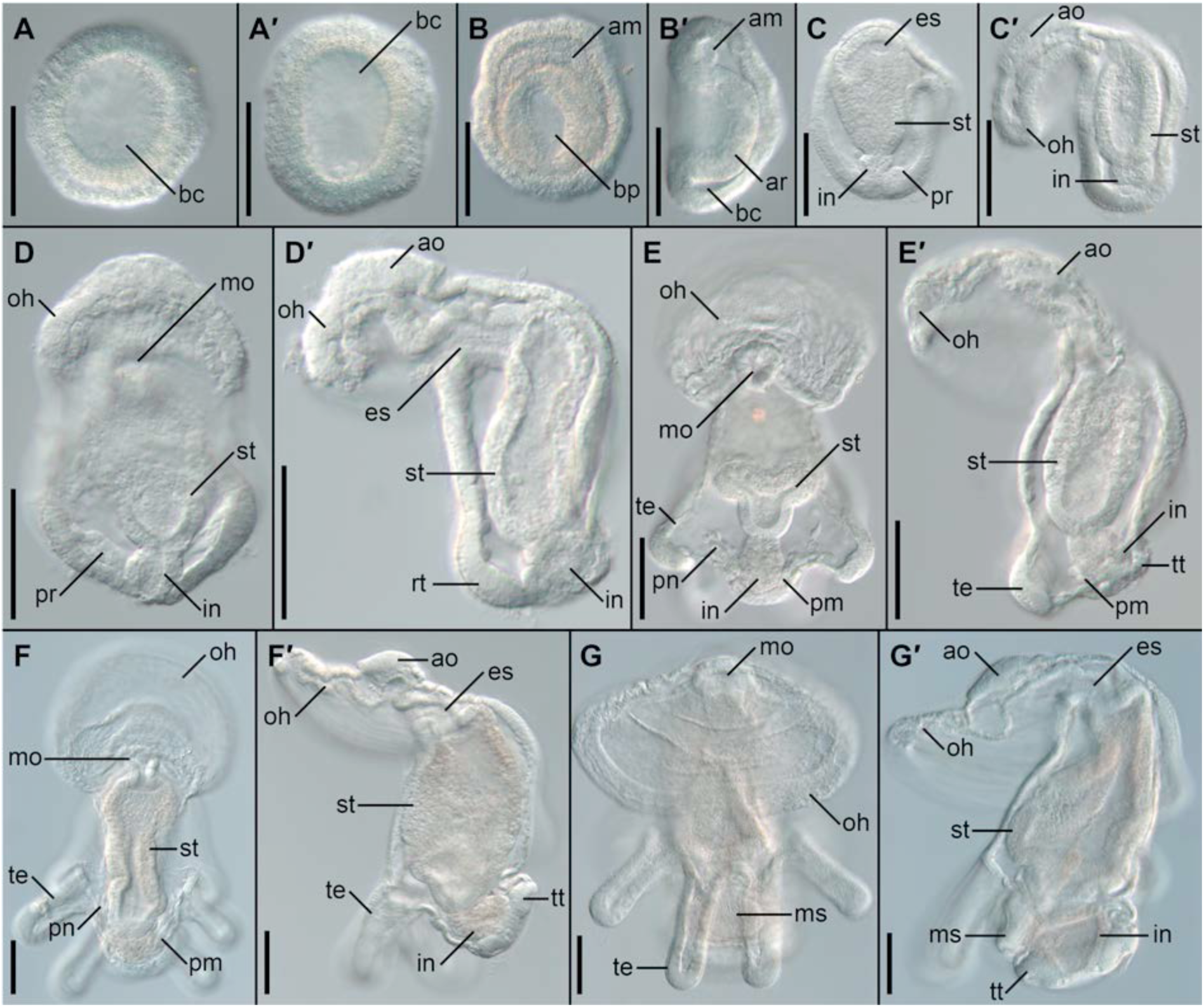
Development of *Phoronopsis harmeri*. Blastula, 12 hpf (A, A’); gastrula, 24 hpf (B, B’); early larva, 42 hpf (C, C’); preactinotrocha, 56 hpf (D, D’); actinotrochae: 3 dpf (E, E’), 5 dpf (F, F’) and 7 dpf (G, G’). For each developmental stage the left panel shows embryo or larvae in dorso-ventral view and right panel (marked as’) in lateral view with ventral to the left; anterior is to the top on all panels. Scalebars 50μm. Abbreviations: *am* anterior mesoderm, *ao* apical organ, *ar* archenteron, *bc* blastocoel, *bp* blastopore, *es* esophagus, *in* intestine, *mo* mouth opening, *ms* metasomal sac, *oh* oral hood, *pm* posterior mesoderm, *pn* protonephridium, *pr* protonephridial rudiment, *rt* tentacle rudiment, *st* stomach, *te* tentacle, *tt* telotroch.

### Hox gene expression

We did not detect expression of any of the Hox genes in blastula and gastrula stages (Additional File 1: Fig. S1). Expression of the anterior Hox gene *lab* is detected for the first time during development at the late 6-tentacle actinotrochae stage (Fig. 4A g and h). The gene is expressed in the ventro-posterior ectodermal domain, between the tentacle bases and the telotroch (black arrowhead, Fig. 4A g and h) and in the paired domains of the dorso-lateral posterior mesoderm (red arrowheads, Fig. 4A g and h). Both of the expression domains persist to the 8-tentacle actinotrocha stage (Fig. 4A i and j). At this developmental stage the ectodermal domain is part of the metasomal sac, where *lab* is expressed in the cells of the anterior and bottom portion of the sac (Fig. 5A and A’).

**Fig. 4.**
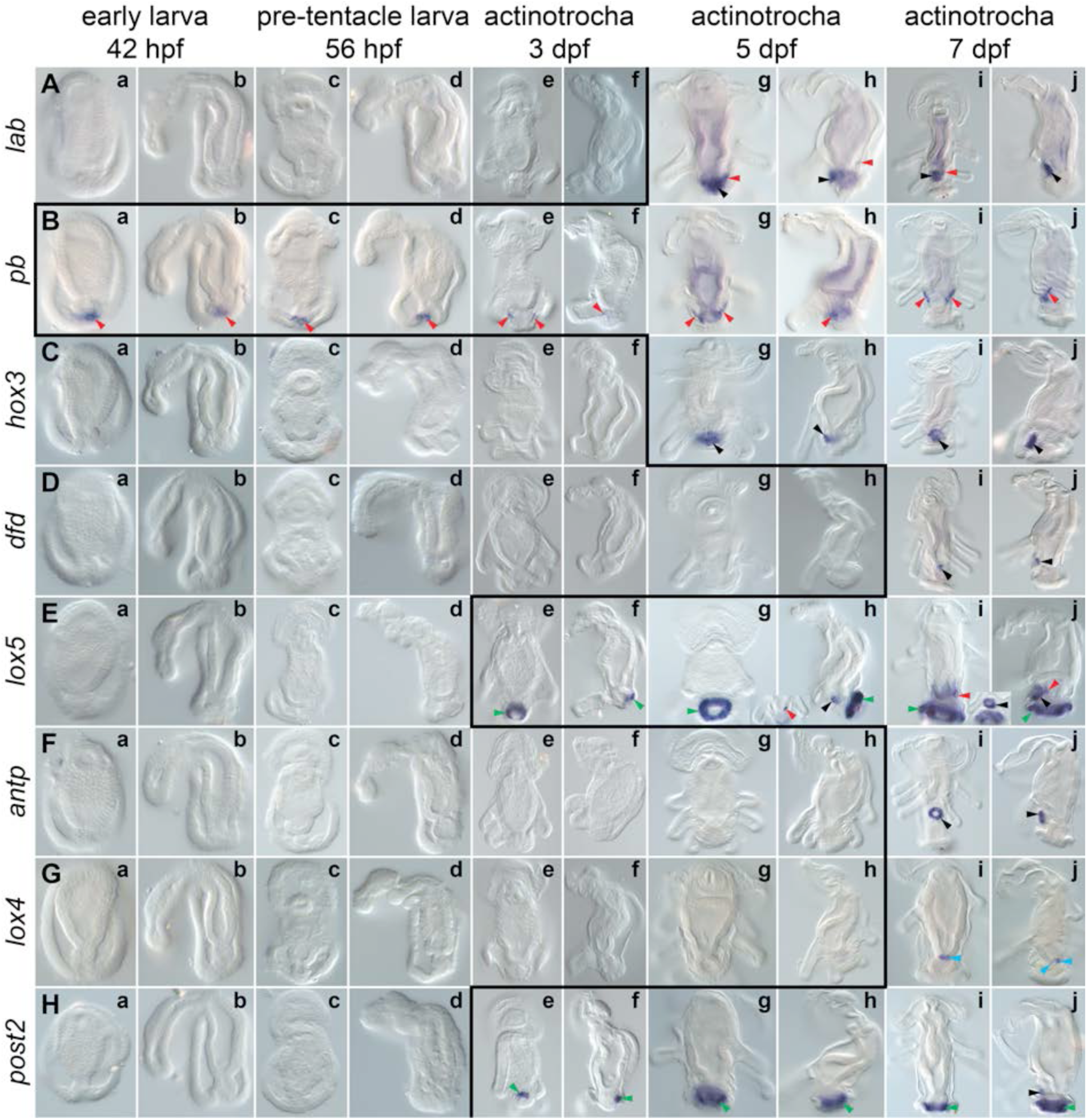
Whole-mount *in situ* hybridization of each Hox gene during larval development of *Phoronopsis harmeri*. Name of each hybridized gene is shown on the left, while developmental stages are indicated on the top. All the stages are presented with anterior to the top. Larvae on panels *a, c, e, g* and *i* are in dorso-ventral view, whereas larvae on panels *b, d, f, h* and *j* in lateral view with ventral to the left. The black line indicates the onset of expression of each Hox gene based on in situ hybridization data. Black arrowheads indicate expression in the metasomal sac, red arrowheads expression in the mesoderm, green arrowheads expression in the telotroch and blue arrowheads expression in the endoderm. The detailed expression patterns are described in the text. Photographs are not to scale.

**Fig. 5.**
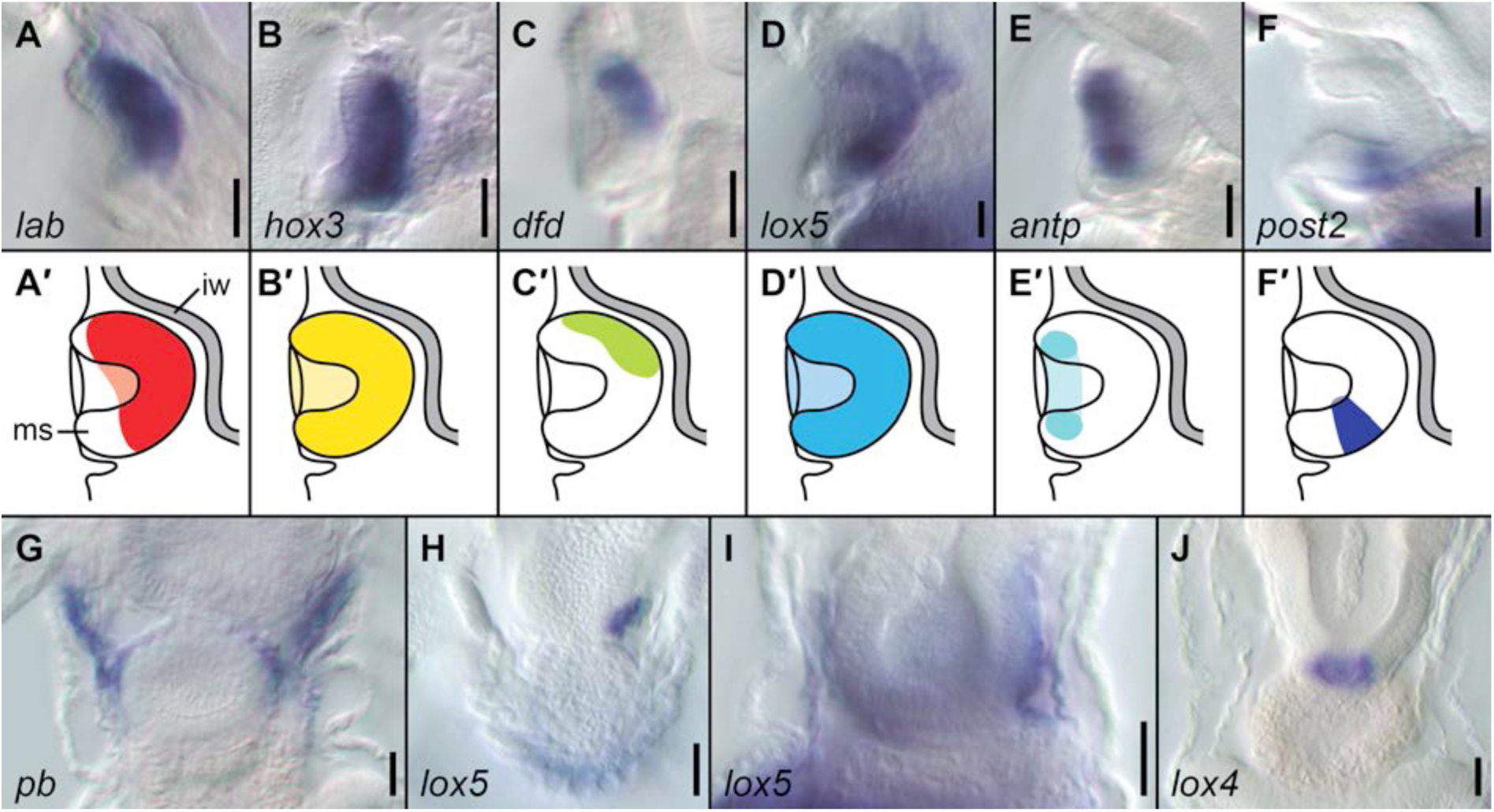
Details of the expression of some of the Hox genes in the actinotrocha larvae of *Phoronopsis harmeri*. Expression of the Hox genes in the metasomal sac of 8-tentacle actinotrochae (A–F) and schematic interpretation of those expression patterns (A’–F’). Expression of *pb* in the telotroch flexors of 8-tentacle actinotrocha (B). Expression of *lox5* in the left mesoderm of 6-tentacle (C) and 8-tentacle actinotrocha (D). Expression of *lox4* in the endoderm of 8-tentacle actinotrocha (E). Scale bars 25μm. Abbreviations: *ms* metasomal sac, *iw* intestinal wall.

The second anterior Hox gene, *pb*, is the earliest expressed among all Hox genes in *P. harmeri* as its expression can be already detected in the early larva stage (42 hpf) in the cells at the border of intestine and anus (red arrowheads, Fig. 4B a and b). This expression domain remains in the pre-tentacle stage (56 hpf, 4B c and d) and later it is restricted to the small lateral domains (Fig. 4B e and f), which in late 6-tentacle actinotrochae expands and the gene labels lateral, mesodermal sheet-like structures, extending along posterior ectoderm (red arrowheads Fig. 4B g and h). In 8-tentacle actinotrochae *pb* is expressed in two mesodermal stripes, which extends from the metasomal sac to the base of tentacles (Fig. 4B i and j, 5G) and corresponds to the telotroch flexor muscles, as described in the late actinotrocha of *P. harmeri* based on phalloidin staining(81).

*Hox3* expression is detected in the late 6-tentacle actinotrochae in an ectodermal domain between the tentacle bases and telotroch (black arrowhead, Fig. 4C g and h). At the 8-tentacle actinotrocha stage *hox3* is uniformly and exclusively expressed in the ectodermal cells of metasomal sac (black arrowheads, Fig. 4C i and j; Fig 5B and B’).

*Dfd* expression initiates only at the 8-tentacle actinotrocha stage (Fig. 4D i and j), where the gene is expressed in a small, anterior subset of the metasomal sac (Fig. 5C and C’).

Transcripts of the gene *lox5* are detected first in the early 6-tentacle actinotrocha in posterior cells of the developing telotroch (green arrowhead, Fig. 4E e and f). Later on, *lox5* remains expressed in the telotroch, expanding its expression domain to the entire structure (green arrowheads, Fig. 4E g–j). Two additional expression domains of *lox5* also appear: the metasomal sac rudiment (black arrowhead, Fig. 4E h), which later encompass the entire metasomal sac (black arrowheads Fig. 4E j and inset between i and j; Fig 5D and D’), and an asymmetric domain in the left ventro-lateral posterior mesoderm, located between metasomal sac, midgut and left body wall (red arrowheads Fig. 4 i and j and inset between g and h; Fig. 5H, I).

Expression of *antp* is not detected until the 8-tentacle actinotrocha stage. Transcripts of the gene are found in ectodermal cells around the opening of the metasomal sac (black arrowheads, Fig. 4F i and j; Fig. 5E and E’), which in a dorso-ventral view look like a ring on the ventral body surface between the tentacles base and the telotroch (Fig. 4F i).

Similarly, *lox4* expression is not detected until the 8-tentacle actinotocha stage, where the gene is exclusively labeling the ring of the endodermal cells at the junction between stomach and intestine (blue arrowheads, Fig. 4G i and j; Fig. J).

The only posterior Hox gene, *post2*, is expressed from the early 6-tentacle actinotrocha (3dpf) in the telotroch (green arrowheads, Fig. 4H e and f), initially in the posterior portion of the organ but later on the expression domain is uniformly surrounding the anus (green arrowheads, Fig. 4H g–j). However, compared to *lox5* expression (which is also demarcating the telotroch), *post2* labels only the inner ring of epidermal cells of the organ (compare Fig. 4E g–j and 4H g–j) and not the entire structure. At the 8-tentacle actinotrocha stage the gene *post2* is additionally expressed in the small posterior portion of the metasomal sac (black arrowhead, Fig. 4H j; Fig 5F and F’).

## Discussion

### Hox gene complement in Phoronida

Similar to the results of the investigation of *Ph. australis* genome, we identified 8 Hox genes in *P. harmeri*, which represent single copies of conserved orthologues of the spiralian Hox genes (Fig. 1A, 2). Luo et al.(89) reported that *Ph. australis* lacks *scr* and *post1* orthologues and suggested that those two genes, which are expressed in the shell forming tissues of brachiopods(16, 40), might be lost in Phoronida due to the evolutionary reduction of the shell in this clade. Accordingly, we also did not identify orthologues of *scr* and *post1* in *P. harmeri*, strengthening the idea that those two Hox genes were absent in the common ancestor of all phoronids. Recent studies are favoring a sister group relationship between phoronids and ectoprocts(52-55, 94), the latter of which possesses a mineralized external shell similarly as brachiopods. However, the Hox gene survey using degenerate polymerase chain reaction primers in ectoproct *Crisularia* (*Bugula*) *turrita* did not retrieve neither *scr* nor *post1* sequences(95), which questions the possible correlation between loss of *scr* and *post1* and the reduction of shell secreting tissues in phoronid lineage. Additionally, the proposed loss of shell in Phoronida, taking into account their currently established phylogenetic position, is disputable itself. Parsimonious reconstruction of the presence of such structure in the last common ancestor of Lophophorata is ambiguous and depends solely on the usage of accelerated vs. delayed transformation algorithm for character optimization, the choice of which is purely arbitral(96).

The gene that was identified as *lox2* by Lou et al.(89) in the genome of *Ph. australis* (and its orthologue in *P. harmeri*) was recovered in our gene orthology analysis as orthologue of *antp* (Fig. 2). Inspection of the phylogenetic tree available in Lou et al. shows that the assessment of the orthology of this gene was tentative, since the gene was actually placed outside of the well-defined clade of *lox2* in their analysis(89). Identification of this gene as *antp* instead of *lox2* is further supported by its position in the genome of *Ph. australis*, which corresponds to the *antp* position in the spiralian species with conserved, organised Hox cluster (Fig. 1A). Additionally, alignment of those phoronid genes with *antp* and *lox2* shows that they lack typical signatures of *lox2*(93) and instead are more similar to the *antp* sequence (Additional File 1: Fig. S2). Consequently, both phoronid species lack an orthologue of *lox2*, the absence, which is apparently shared by Phoronida with other Lophophorata(16, 89, 90, 95) as well as with some other Spiralia – i.e. Rotifera(34, 97) and Platyhelminthes(42, 98). *Lox2* was originally described from leeches(99, 100) to be later proposed as an evolutionary innovation of Lophotrochozoa ((93), sensu = Spiralia(101)). However, its orthologues are so far identified only in annelids (e.g. (27, 46, 91, 93, 99, 100, 102)), nemerteans(89), molluscs (e.g. (30, 36, 41, 91, 93, 103-106)) and possibly kamptozoans(107) (however, in the latter the *lox2*-like sequence lacks most of the residues considered as *lox2* signature, Additional File 1: Fig. S2). This indicates that *lox2* evolved only after split of the common ancestor of those clades from remaining Spiralia and does not belong to the ancestral hox complement of all Spiralia(16). Whether absence of *lox2* in lophophorates is plesiomorphic or represents evolutionary reversal depends on the position of Lophophorata within Spiralia, which is still debatable and not fully resolved(52-55, 94).

### Hox genes in Phoronida do not show traces of collinear expression

When assuming the presence of a similar gene order in the Hox cluster of *P. harmeri* as in *Ph. australis* then the former does not show any traces of temporally or spatially collinear expression of Hox genes. This is especially intriguing taking into account that *Ph. australis* has highly organized Hox cluster and collinear expression (especially in its temporal aspect) has been proposed as a main evolutionary factor responsible for conservation of organization of Hox cluster(9, 11-16, 49). Therefore, either another mechanism is responsible for Hox cluster conservation in Phoronida or the two discussed phoronid species varies greatly in the cluster organization and/or Hox gene expression patterns.

6 out of 8 identified Hox genes are expressed in the metasomal sac (*pb* and *lox4* being the only two, which expression was not detected in the structure) and already at the stage of 8-tentacle actinotrocha some of those genes (*lab, dfd, antp, post2*) show differentiated expression in a particular region of the sac (Fig. 5A’), although without any clear pattern along the future A-P axis. However, it is possible that in the competent larvae (when the metasomal sac is a fully formed, elongated structure; (81, 82)), the expression of particular Hox genes is restricted to the different regions of the trunk rudiment and shows some traces of staggered expression along the future A-P axis of the worm body. Hence, the future investigation of the Hox expression in competent larvae and freshly metamorphosed juveniles can reveal spatial collinearity obliterated in the early stages of metasomal sac development or eventually confirm a lack of collinear Hox expression throughout entire development of phoronids.

### Germ layer-specific expression of Hox genes in Spiralia

Although Hox genes in Bilateria are predominantly expressed in the ectoderm (including nervous system) and their ectodermal expression is often considered as an ancestral feature(14, 28, 34), in various spiralian species certain Hox genes are also expressed in mesoderm, endoderm and clade-specific structures like chaetal sacs or shell fields (e.g. (16, 23, 24, 27, 29, 31, 35, 36, 39-41, 46); Tab. 1). Inclusion of the data on Hox expression in Phoronida gives some new insight into the understanding of the evolution of germ-layer specific Hox expression in Spiralia. *P. harmeri*, similarly as two investigated brachiopod species(16, 40), seems to lack expression of any of the Hox genes in the nervous system, the peculiarity that might actually represent apomorphy of Lophophorata. Three of the Hox genes – *pb, hox3* and *dfd* – were shown to be differentially expressed along the A-P axis in the mesoderm of brachiopod larvae(16). Out of those three genes, only *pb* (which mesodermal expression is actually lacking in craniiformean *Novocrania anomala*(16)) is expressed mesodermally in *P. harmeri*, indicating that cooption of *hox3* and *dfd* into mesoderm patterning occurred after the split of brachiopods and phoronids. Interestingly, the expression of *pb* shows a remarkable similarity between *P. harmeri* and the rynchonelliformean brachiopod *Terebratalia transversa*. In both species the gene is initially expressed in a broader mesodermal domain, while later its expression is restricted to the single pair of muscles (telotroch flexors in *P. harmeri* and anterior shell adductor muscles in *T. transversa*(40)). Taking into account that in advanced phoronid larvae *pb*-positive cells are exclusively mesodermal, the cells at the border between hindgut and anus which express *pb* in early larval stages of *P. harmeri* might represent early precursors of the posterior mesoderm. This corresponds to the morphological observation that posterior mesoderm originates in early larvae from the posterior part of digestive tract after formation of protonephridial rudiments(76). Comparison of Hox gene expression across Spiralia (Tab. 1) allows the observation that *pb* is mesodermally expressed in many species and it is likely that mesodermal expression of *pb* represents an ancestral condition in Lophotrochozoa (*sensu stricto*(101)). Last but not least, the endodermal expression of *lox4* in *P. harmeri* is a peculiar and derived feature as this gene is expressed in other Spiralia in ectoderm, nervous system or mesoderm. In general, among Spiralia, the Hox genes are rarely expressed in the endodermal structures (Tab 1.).

**Table 1.** Expression of Hox genes in spiralian species.

| species | clade | reference | Hox gene |  |  |  |  |  |  |  |  |  |  |
| --- | --- | --- | --- | --- | --- | --- | --- | --- | --- | --- | --- | --- | --- |
|  |  |  | Lab | Pb | Hox3 | Dfd | Scr | Lox5 | Antp | Lox4 | Lox2 | Post2 | Post1 |
| *Phoronopsis harmeri* | Phoronida | this study | ectoderm,<br>mesoderm | mesoderm | ectoderm | ectoderm | gene absent | ectoderm,<br>mesoderm | ectoderm | endoderm | gene absent | ectoderm | gene absent |
| *Terebratalia transversa* | Brachiopoda | (16, 40) | chaetal sac | ectoderm,<br>mesoderm | ectoderm,<br>mesoderm | ectoderm,<br>mesoderm | shell field | ectoderm | ectoderm | ectoderm | gene absent | ectoderm | chaetal sac |
| *Novocrania anomala* | Brachiopoda | (16) | chaetal sac | ectoderm | ectoderm,<br>mesoderm | ectoderm,<br>mesoderm | shell field | ectoderm | ectoderm | unknown | gene absent | unknown | gene absent |
| *Capitella teleta* | Annelida | (46) | ectoderm,<br>nervous<br>system | ectoderm,<br>nervous<br>system | ectoderm,<br>nervous<br>system | ectoderm,<br>nervous<br>system | ectoderm,<br>nervous<br>system | ectoderm,<br>nervous<br>system | ectoderm,<br>nervous<br>system | ectoderm,<br>nervous<br>system | ectoderm,<br>nervous<br>system | ectoderm,<br>nervous<br>system | chaetal sac |
| *Alitta virens* | Annelida | (27, 31) | ectoderm,<br>chaetal sac,<br>nervous<br>system | ectoderm,<br>mesoderm | ectoderm | ectoderm,<br>nervous<br>system | ectoderm,<br>nervous<br>system | ectoderm,<br>nervous<br>system | ectoderm,<br>nervous<br>system | ectoderm,<br>nervous<br>system,<br>mesoderm | ectoderm,<br>nervous<br>system | ectoderm,<br>nervous<br>system,<br>mesoderm,<br>endoderm | chaetal sac |
| *Chaetopterus variopedatus* | Annelida | (23) | ectoderm,<br>nervous<br>system | ectoderm,<br>nervous<br>system,<br>mesoderm | ectoderm,<br>nervous<br>system | ectoderm,<br>nervous<br>system | ectoderm,<br>nervous<br>system | unknown | unknown | unknown | unknown | unknown | unknown |
| *Acanthochitona crinita* | Mollusca | (35, 36) | ectoderm,<br>mesoderm | mesoderm | mesoderm | ectoderm,<br>mesoderm | mesoderm | ectoderm,<br>mesoderm | mesoderm | ectoderm,<br>mesoderm | mesoderm | ectoderm,<br>mesoderm | gene absent |
| *Antalis entalis* | Mollusca | (39) | shell field,<br>nervous<br>system | ectoderm,<br>nervous<br>system | ectoderm,<br>shell field,<br>nervous<br>system,<br>mesoderm | nervous<br>system,<br>mesoderm | nervous<br>system,<br>mesoderm | ectoderm,<br>nervous<br>system | gene absent | ectoderm,<br>nervous<br>system,<br>mesoderm | unknown | ectoderm,<br>nervous<br>system | ectoderm,<br>shell field,<br>nervous<br>system |
| *Lottia goschimai* | Mollusca | (41) | shell field,<br>nervous<br>system | shell field,<br>nervous<br>system | shell field,<br>nervous<br>system | shell field,<br>nervous<br>system | shell field,<br>nervous<br>system | shell field,<br>nervous<br>system | nervous<br>system | shell field,<br>nervous<br>system | ectoderm,<br>shell field,<br>nervous<br>system | shell field,<br>nervous<br>system | nervous<br>system |
| *Brachionus manjavacas* | Rotifera | (34) | gene absent<br>(?) | nervous<br>system | nervous<br>system | nervous<br>system | unknown | nervous<br>system | gene absent<br>(?) | gene absent<br>(?) | gene absent<br>(?) | gene absent<br>(?) | gene absent<br>(?) |

### Hox gene expression and the nature of actinotrocha larvae

We showed that in *P. harmeri* Hox genes are not expressed during embryogenesis, when the larval body is formed, but instead they are expressed in the metasomal sac (which will contribute to the adult trunk epidermis), posterior mesoderm (which contributes to the mesodermal structures in the adult trunk), the small posterior portion of the endoderm (which during metamorphosis descent into the trunk rudiment forming loop of the U-shaped intestine) and the telotroch. In many of the investigated Bilateria, Hox genes are expressed already during early developmental stages and, if the biphasic life cycle is present, they are involved in the formation of both larval and adult body plans (e.g. (16, 27, 29-31, 40, 41, 45, 46, 48)). However, there are some animals that, similarly as phoronids, deviate from this general pattern. For instance, in pilidiophoran nemerteans(37) as well as in indirectly developing sea urchins(21, 22, 47) and hemichordates(38), the larvae develop without expressing Hox genes (or with the expression of just few of them). On the other hand, in ascidian tadpole larva, Hox genes pattern the developing larval nervous system, but only two posterior Hox genes are expressed after metamorphosis during development of juvenile animal(44).

Two evolutionary processes have been proposed as responsible for those observed differences in the Hox gene expression during formation of the larval and adult bodies. According to the first hypothesis, an ancestral body plan – the adult worm – retains patterning by Hox genes, while the new form is intercalated into to the life cycle, and uses another molecular mechanism to provide primary positional information to the cells of developing body(37, 45). Another concept was proposed to explain phenomenon observed during larval development of a hemichordate *Schizocardium californicum*(38, 92). Although metamorphosis in this species is not so drastic(108), the larvae develops also without expression of any Hox genes, which instead are expressed only late during larval development and only in the posterior region of the competent larvae, from which the trunk of the juvenile worm will develop during metamorphosis(38). In the so-called “head larva”-hypothesis it was proposed that larval body represents the homologue of only the head region of the future animal while the trunk is added later during post-larval development(38). The “head larva” develops without expression of the Hox genes, which often are involved only in the patterning of the trunk but not the head(38, 45, 89, 109, 110). Hox gene expression becomes activated only after the onset of trunk development and that is why Hox genes pattern only the adult body(38).

Both of those hypotheses might be applied to explain the Hox expression patterns we observed in *P. harmeri*. According to the first hypothesis, the actinotrocha larva might represent an evolutionary novelty in the life cycle of phoronids, which was intercalated in the phoronid lineage and that is why it is not patterned by an ancestral Hox gene system. Such idea is supported by the fact, that the actinotrocha does not bear obvious homology to any other spiralian larvae(80, 111, 112). Moreover, the actinotrocha is lacking in *Ph. ovalis*(60), which is the sister species to all remaining phoronids(62-64), suggesting that the actinotrocha was not even present in the most recent ancestor of all Phoronida, but instead appeared during the evolution after split between *Ph. ovalis* and the remaining phoronids. On the other hand, from the morphological point of view, the tentacles of actinotrocha larvae are homologous to the tentacles of the lophophore in the adult worm ((73, 82, 112); Fig. 1C), and the adult lophophore exhibits the molecular signature of a bilaterian head(89). As tentacles are positioned posteriorly in the early actinotrocha, one can conclude that on a morphological basis the actinotrocha is mostly composed of the head region. Following such interpretation, all of the Hox genes are expressed in the structures, which will contribute to the adult trunk tissues but are not expressed in the developing future head (and hence in the most of the larval body). Accordingly, based on a body region specific transcriptome, it has been demonstrated that in adults of *Ph. australis* Hox genes are not expressed in the lophophore, while their expression is detectable in the trunk and posterior ampulla(89). Similarly, in rhynchonelliformean and craniiformean brachiopods none of the Hox genes are expressed in the larval anterior lobe(16, 40), which contributes to the lophophore after metamorphosis(112). A lack of Hox expression in the adult lophophore tissue (as opposed to the remaining body regions) was shown for the linguliformean *Lingula anatina*, based on the tissue specific transcriptomics(89). Additionally, our study shows that two of the Hox genes (*lox5* and *post2*) are expressed in the telotroch, which represent a truly larval structure, that is lost during metamorphosis(73, 82), therefore Hox genes are indeed, albeit to only a limited degree, involved in the larval development. Hox gene expression in the larval telotroch can be explained by the fact that telotroch represents a truly “posterior” structure, belonging to the post-head body region even in the earliest, “head dominated” actinotrocha. The “head larva” interpretation is additionally in agreement with the recent investigation of the expression of anterior, posterior and endomesodermal molecular markers during embryonic and early larval development of *P. harmeri*(85). Andrikou et al.(85) have demonstrated that although some of the markers have peculiar expression patterns, most of them show conserved expression between phoronids and various Spiralia, especially rhynchonelliformean brachiopods. This indicates that the A-P axis and endomesodermal tissues of phoronid larvae are, to the certain degree, specified similarly as in other Spiralia(85). Additionally, the gene *six3/6*, which is considered as marker of anterior, hox-free head region (38, 45, 89, 109, 110, 113) is expressed in *P. harmeri* not only in the apical organ but also in certain more posteriorly located domains, including stomach and ventral ectoderm(85).

It is, however, important to stress that the hypotheses of the “larval intercalation” and “head larva” are not mutually exclusive. It is possible that the actinotrocha was intercalated into life cycle of Phoronida after split of *Ph. ovalis* lineage and that it originated from the precocious development of the head structures or delayed development of the adult trunk.

### Conclusions

We identified 8 Hox genes (*lab, pb, hox3, dfd, lox5, antp, lox4* and *post2*) in the transcriptome of *Phoronopsis harmeri* showing that Hox complement in this species is the same as in *Phoronis australis*, which genome, including organized Hox cluster, was recently described. Both phoronid species lack genes *scr* and *post1*, which likely belong to the ancestral Hox cluster of Spiralia. Phoronids, similarly as closely related brachiopods and bryozoans, as well as some other Spiralia, lack gene *lox2*. This absence indicates that *lox2* evolved in the common ancestor of annelids, nemerteans molluscs and possibly entoprocts, while its lack is plesiomorphic among Spiralia.

Hox gene expression is activated late during the development of *P. harmeri*. The larval body develops without expressing any of the Hox genes, which instead are expressed in the tissues of the prospective rudiment of the adult worm (metasomal sac, hindgut and posterior mesoderm) and in the telotroch. Such expression might result either from the intercalation of actinotrocha larva into the ancestral life cycle of phoronids or from the fact that the early larva of phoronids represents a “head larva”, composed mostly of the anterior head structures, which develops without expressing any Hox genes. Those two explanations are not mutually exclusive and we propose that a new body form was intercalated to the phoronid life cycle by precocious development of the anterior structures or by delayed development of the trunk rudiment in the ancestral phoronid larva. Such hypothesis can be tested in future by investigation of the head specific genes (e.g. *foxG1, pax6, lim1/5* or *soxB*(38, 89, 109, 114, 115)) during embryonic, larval and post-metamorphic development of *P. harmeri*. Additionally, study of the Hox gene expression during the development of *Phoronis ovalis*, a sister group to all remaining Phoronida, which lacks the actinotrocha larva stage and develops through a creeping, worm-like larva, might be also used to test our hypothesis.

## Methods

### Animal collection and fixation

Gravid females of *P. harmeri* Pixell, 1912 were collected in Bodega Bay (38° 18’ 51.9012” N 123° 3’ 12.3012” W) in California during April and May. The animals were opened in the laboratory and eggs (fertilized during dissection by sperm stored in the coelom of females) were transferred to the clean cultures with filtered see water (as described in e.g. (78, 84, 85)). Embryos are initially lecitotrophic, but after formation of the gut larvae requires feeding, hence concentrated *Rhodomonas* or *Rhinomonas* algae were added to the cultures. Water in the larval cultures was exchange every 2-3 days, followed by the addition of fresh algae. Embryos and larvae on desired developmental stages were relaxed with 8% MgCl_2_, fixed in 3.7% formaldehyde and subsequently washed in phosphate buffer with 0.1% Tween-20. Fixed animals were stored in 100% methanol in -20°C.

### Hox genes identification and orthology assessment

We searched transcriptome of *P. harmeri* with reciprocal TBLASTN using 8 Hox protein sequences from *Phoronis australis*. The top 10 homeodomain-containing BLAST hits from each search were blasted back to the protein database at NCBI (http://blast.ncbi.nlm.nih.gov/) and if any Hox gene was a top reciprocal hit, the sequence was considered to be a putative Hox gene. We identified 8 sequences, which passed this reciprocal test and translated them to the proteins using CLC Main Workbench 7. Orthology of particular phoronid Hox genes was assessed based on the results of phylogenetic analysis. In order to construct alignment, amino acid sequences of Hox transcription factors and nucleotide sequences of Hox genes from several spiralian species were obtained from GenBank (https://www.ncbi.nlm.nih.gov/genbank/), ENSEMBL genome data base (https://www.ensembl.org/index.html) and website of Marine Genomics Unit of Okinawa Institute of Science and Technology (http://marinegenomics.oist.jp). For the nucleotide sequences ORF were determined based on BLAST results at NCBI and sequences were translated into proteins using CLC Main Workbench 7. All spiralian sequences used in this study with their source and accession number are provided in the Additional File 2: Table S1. The spiralian Hox protein sequences, including putative Hox genes of *P. harmeri*, were aligned in CLC Main Workbench 7 and then the alignment was manually trimmed to contain the conserved homeodomain (60 aa), five amino acids 5' of the homeodomain, and eight aa 3’ of the homeodomain (the trimmed alignment in FASTA format is available in the Additional File 2). ProtTest3 (116) was used to determine the best-fitting substitution model (JTT+I+G). Bayesian analysis was run in MrBayes v3.2.6 (117, 118) with the JTT+I+G substitution model in two independent runs, each with 4 Marcov chains (three heated and one cold) with 3.000.000 generations sampled every 500 generations. The Gsx sequence of *Capitella teleta* was designated as outgroup. The first 25% of samples were discarded as burn-in and the remaining trees were used to calculate posterior probability values and construct consensus tree, which was visualized and adjusted in FigTree v1.4.3.

All new sequences obtained and identified in this study were uploaded to the GenBank (accession numbers MN443105–MN443114).

### Gene cloning and probe synthesis

Fragments of each Hox gene were amplified from cDNA libraries from mixed larval and adult tissues using gene specific primers (provided in Additional file2: Table S2) designed in MacVector 11.0.4 based on the sequences found in the transcriptome. PCR products were cloned into pGEM-T Easy vectors (Promega, USA) and then transformed into competent *Escherichia coli* cells. Plasmid DNA was isolated and sequenced in both forward and reverse directions using T7 and SP6 primers. Labeled antisense RNA probes were transcribed from linearized DNA using digoxigenin-11-UTP (Roche, USA) according to the manufacturer’s instructions.

### In situ hybridization and light microscopy

Single whole-mount in situ hybridization was performed following an established protocol(119) with proteinase K digestion time of 2 minutes. Probes were hybridized at a concentration of 1ng/μl at 67°C for approximately 72 h, detected with anti-digoxigenin-AP antibody in 1:5000 concentration in blocking buffer and visualized with nitroblue tetrazolium chloride and 5-bromo-4-chloro-3-indolyl phosphate. Embryos and larvae were mounted in 70% glycerol and examined with Zeiss Axiocam HRc connected to a Zeiss Axioscope Ax10 using bright-field Nomarski optics.

### Image processing and figure preparation

Light micrographs were adjusted in Adobe Photoshop CS6 for contrast and assembled in Adobe Illustrator CS6. All figures were prepared in Adobe Illustrator CS6.

## Declarations

### Ethics approval and consent to participate

Studies of phoronids do not require ethics approval or consent to participate.

### Consent for publication

Not applicable.

### Availability of data and material

Sequences generated and analyzed in this study have been deposited in NCBI’s GenBank database under accession numbers MN443105–MN443114. All remaining data generated or analyzed during this study are included in this published article.

### Authors’ contributions

Both authors conceived the study, analyzed data and contributed to writing. LG took part in sample collection, conducted gene search and orthology assessments, performed in situ hybridization, arranged figures and drafted the manuscript. Both authors read and approved the final manuscript.

## Acknowledgements

We thank Yale Passamaneck who contributed with phoronid collection and spawning. Chris Lowe from Hopkins Marine Station of Stanford University is greatly acknowledged for hosting LG in his lab in April – May 2018 and for sharing a transcriptome of *P. harmeri* before it was made public. We also would like to thank Carmen Andrikou and Andrea Orús Alcalde as well as Chris Lowe and Paul Bump for contributing with animal collection in April – May 2018. Carmen Andrikou is additionally greatly acknowledged for reading and commenting on the manuscript.

