## Supplementary material for "Hox gene expression during the development of the phoronid *Phoronopsis harmeri*": Figure S1 and Figure S2

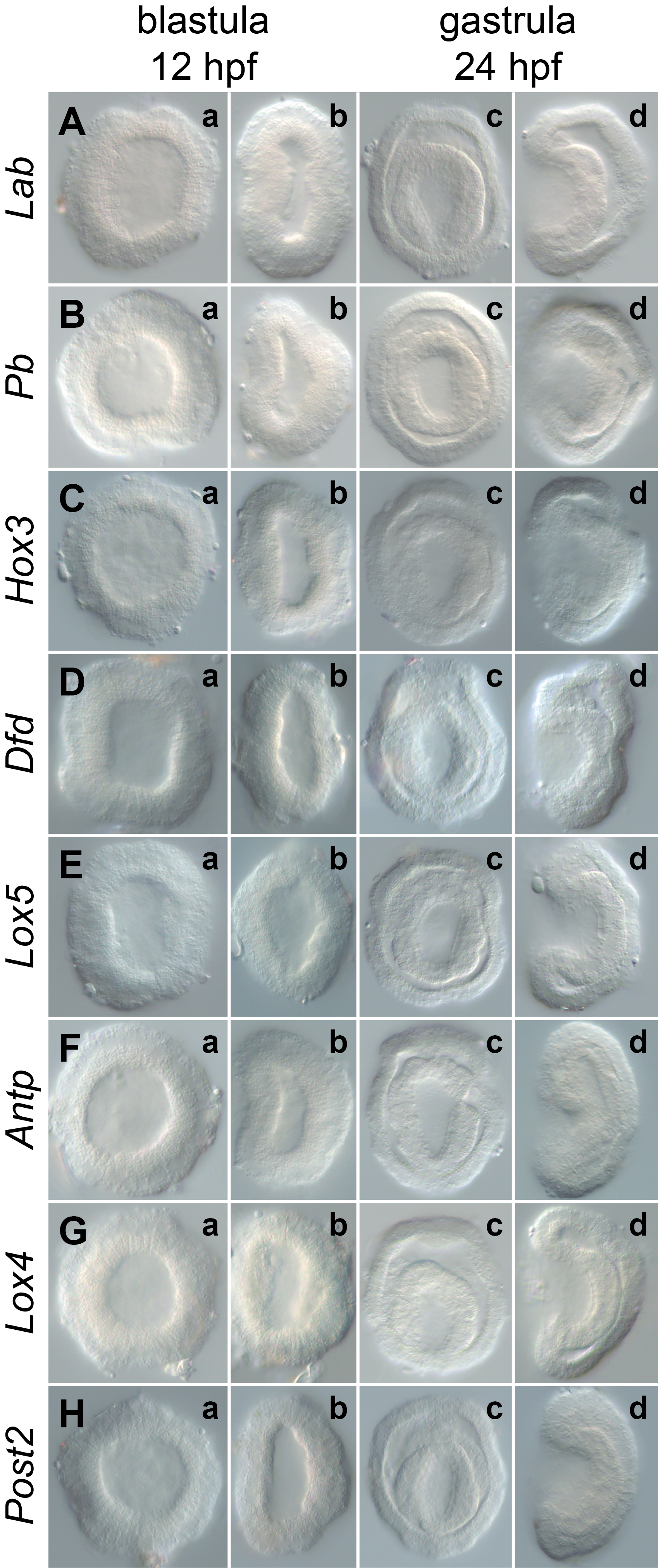

**Figure S1.** Lack of Hox gene expression in early developmental stages of *P. harmeri*. Name of each hybridized gene is shown on the left, while developmental stages are indicated on the top. Embryos on panels a and c are in the vegetal view, whereas embryos on panels b and d in the lateral view. Anterior is to the top on all panels. Photographs are not to scale.

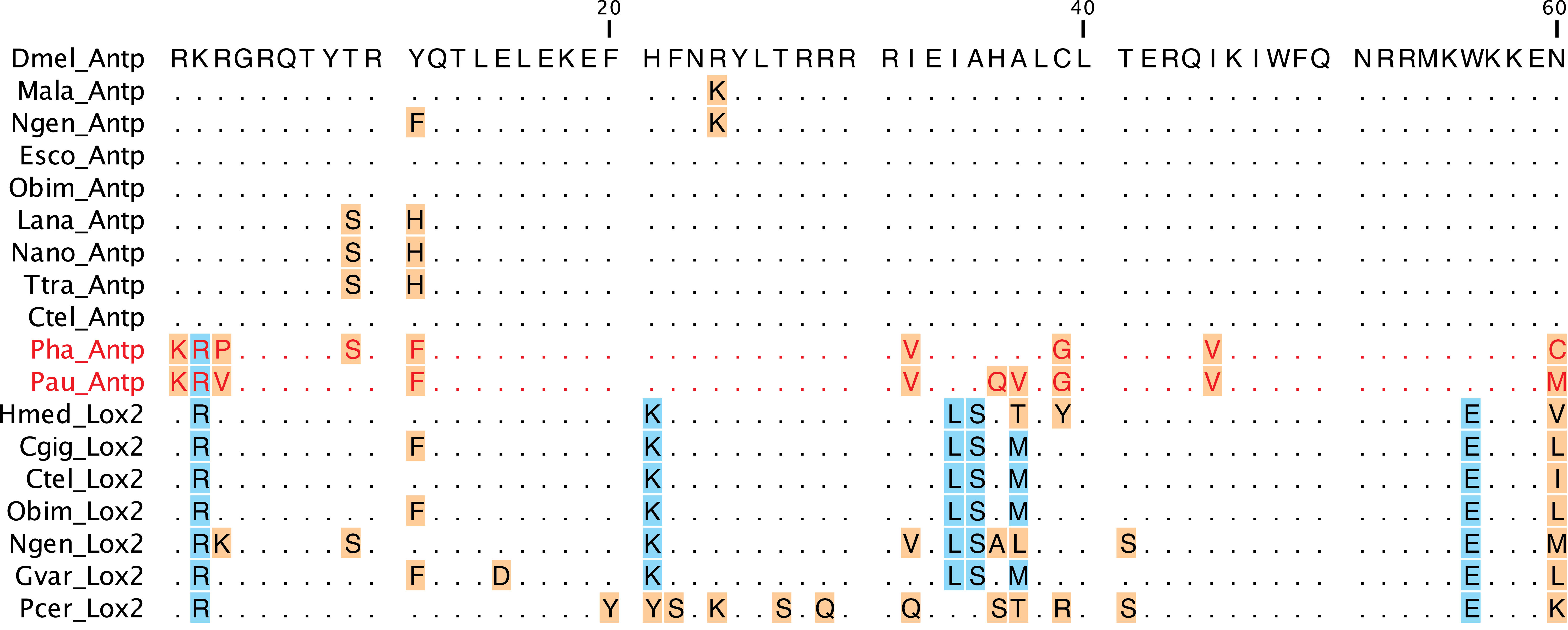

**Figure S2.** Sequences of homeodomains of spiralian *antp* and *lox2* aligned with the sequence of *Drosophila melanogaster* *antp*. Note that although sequences identified herein as phoronid *antp* share one amino acid with *lox2*, they lack remaining 5 signatures typical for *lox2*. Dots indicate identity to *D. melanogaster* *antp*, *lox2* specific signatures are highlighted in blue and remaining residues that differ from *D. melanogaster* *antp* are highlighted in orange. Phoronid sequences are in red. Dmel stands for *Drosophila melanogaster*, Hmed for *Hirudo medicinalis*, Pcer for *Piedicellina cernua*. Sequence of *PcerLox2* was obtained from GenBank (accession number KP691980). For remaining species abbreviations and source of sequences see Additional file 2: Tab. S1.
