## Supplementary material for "Hox gene expression during the development of the phoronid *Phoronopsis harmeri*": Table S1 and Table S2

**Table S1.** Sequences used in the phylogenetic assessment of Hox genes orthology.

| **species** | **shortcut** | **clade** | **gene** | **accession no** | **source** |
| --- | --- | --- | --- | --- | --- |
| *Flaccisaggita enflata* | Fenf | Chaetognatha | Hox1 | ABS18809.1 | GenBank |
|  |  |  | Hox3 | ABS18810.1 |  |
|  |  |  | Hox4 | ABS18811.1 |  |
|  |  |  | Hox5 | ABS18812.1 |  |
|  |  |  | Hox6 | ABS18813.1 |  |
|  |  |  | Hox8 | ABS18814.1 |  |
|  |  |  | MedPost | ABS18817.1 |  |
|  |  |  | Posta | ABS18815.1 |  |
|  |  |  | Postb | ABS18816.1 |  |
| *Capitella teleta* | Ctel | Annelida | lab | ABY67952 | GenBank |
|  |  |  | pb | ABY67953 |  |
|  |  |  | Hox3 | ABY67954 |  |
|  |  |  | Dfd | ABY67955 |  |
|  |  |  | Scr | ABY67956 |  |
|  |  |  | Lox5 | ABY67957 |  |
|  |  |  | Antp | ABY67962 |  |
|  |  |  | Lox4 | ABY67958 |  |
|  |  |  | Lox2 | ABY67959 |  |
|  |  |  | Post1 | ABY67961 |  |
|  |  |  | Post2 | ABY67960 |  |
|  |  |  | Gsx | AAZ23124.1 |  |
|  |  |  | Cdx | AAZ95508 |  |
|  |  |  | Xlox | AAZ95509.1 |  |
|  |  |  | Evx | ABG82164 |  |
| *Crassostrea gigas* | Cgig | Mollusca | Hox1 | CGI_10024083 | ENSEMBL |
|  |  |  | Hox2 | CGI_10024086 |  |
|  |  |  | Hox3 | CGI_10024087 |  |
|  |  |  | Hox4 | CGI_10024091 |  |
|  |  |  | Lox2 | CGI_10018592 |  |
|  |  |  | Lox4 | CGI_10026562 |  |
|  |  |  | Lox5 | CGI_10026565 |  |
| *Octopus bimaculoides* | Obim | Mollusca | Hox1 | Ocbimv22030263 | ENSEMBL |
|  |  |  | Scr | Ocbimv22018468 |  |
|  |  |  | Antp | Ocbimv22036189 |  |
|  |  |  | Post1 | Ocbimv22015181 |  |
|  |  |  | Post2 | Ocbimv22031197 |  |
|  |  |  | Lox2 | Ocbimv22033340 |  |
|  |  |  | Lox4 | Ocbimv22009726 |  |
|  |  |  | Lox5 | Ocbimv22010205 |  |
| *Gibbula varia* | Gvar | Mollusca | HoxA | ACX84671.1 | GenBank |
|  |  |  | Hox2 | ADJ18233.1 |  |
|  |  |  | Hox3 | ADJ18232.1 |  |
|  |  |  | Hox4 | ACX84672.1 |  |
|  |  |  | Hox5 | ADJ18234.1 |  |
|  |  |  | Lox5 | ADJ18235.1 |  |
|  |  |  | Hox7 | ADJ18236.1 |  |
|  |  |  | Lox2 | ADJ18238.1 |  |
|  |  |  | Lox4 | ADJ18237.1 |  |
|  |  |  | Post1 | ACX84673.1 |  |
| *Euprymna scolopes* | Esco | Mollusca | lab | AY330184 | GenBank |
|  |  |  | Hox3 | AY330185 |  |
|  |  |  | Scr | AY330186 |  |
|  |  |  | Lox5 | AY330187 |  |
|  |  |  | Antp | AY330188 |  |
|  |  |  | Lox4 | AY330189 |  |
|  |  |  | Post1 | AAL25811.1 |  |
|  |  |  | Post2 | AY330191 |  |
| *Micrura alaskensis* | Mala | Nemertea | lab | KP762174 | GenBank |
|  |  |  | pb | KP762176 |  |
|  |  |  | Hox3 | KP762173 |  |
|  |  |  | Dfd | KP762180 |  |
|  |  |  | Scr | KP762177 |  |
|  |  |  | Lox5 | KP762179 |  |
|  |  |  | Antp | KP762171 |  |
|  |  |  | Lox4 | KP762175 |  |
|  |  |  | Post2 | KP762178 |  |
| *Notospermus geniculatus* | Ngen | Nemertea | labA | g12075.t1 | marinegenomics.oist.jp |
|  |  |  | labB | g24513.t1 |  |
|  |  |  | pbA | g6836.t1 |  |
|  |  |  | pbB | g12074.t1 |  |
|  |  |  | Hox3A | g30654.t1 |  |
|  |  |  | Hox3B | g6837.t1 |  |
|  |  |  | ScrA | g19273.t1 |  |
|  |  |  | ScrB | g35291.t1 |  |
|  |  |  | DfdA | g33684.t1 |  |
|  |  |  | DfdB | g35826.t1 |  |
|  |  |  | Antp | g19270.t1 |  |
|  |  |  | Lox5 | g19271.t1 |  |
|  |  |  | Lox2 | g26697.t1 |  |
|  |  |  | Lox4 | g26698.t1 |  |
|  |  |  | Post2A | g26696.t1 |  |
|  |  |  | Post2B | g16861.t1 |  |
| *Crisularia turrita* | Ctur | Bryozoa | pb | AAS77225 | GenBank |
|  |  |  | Hox3 | AAS77226 |  |
|  |  |  | Dfd-a | AAS77227 |  |
|  |  |  | Dfd-b | AAS77228 |  |
|  |  |  | Lox5 | AAS77229 |  |
|  |  |  | Post2 | AAS77230 |  |
| *Lingula anatina* | Lana | Brachiopoda | lab | g10891 | ENSEMBL |
|  |  |  | pb | g10890 |  |
|  |  |  | Hox3 | g10889 |  |
|  |  |  | Dfd | g10888 |  |
|  |  |  | Scr | g10887 |  |
|  |  |  | Lox5 | g10886 |  |
|  |  |  | Antp | g10892 |  |
|  |  |  | Post1 | g12396 |  |
|  |  |  | Post2 | g12399 |  |
| *Novocrania anomala* | Nana | Brachiopoda | lab | KX372756 | GenBank |
|  |  |  | pb | KX372757 |  |
|  |  |  | Hox3 | KX372758 |  |
|  |  |  | Scr | KX372759 |  |
|  |  |  | Lox5 | KX372760 |  |
|  |  |  | Dfd | KX372769 |  |
|  |  |  | Antp | KX372770 |  |
|  |  |  | Lox4 | KX372773 |  |
|  |  |  | Post2 | KX372774 |  |
| *Terebratalia transversa* | Ttra | Brachiopoda | lab | KX372761 | GenBank |
|  |  |  | pb | KX372762 |  |
|  |  |  | Hox3 | KX372763 |  |
|  |  |  | Dfd | KX372764 |  |
|  |  |  | Scr | KX372765 |  |
|  |  |  | Lox5 | KX372766 |  |
|  |  |  | Lox4 | KX372767 |  |
|  |  |  | Post2 | KX372768 |  |
|  |  |  | Antp | KX372771 |  |
|  |  |  | Post1 | KX372772 |  |
| *Phoronis australis* | Pau | Phoronida | lab | g5412.t1 | marinegenomics.oist.jp |
|  |  |  | pb | g5413.t1 |  |
|  |  |  | Hox3 | g5414.t1 |  |
|  |  |  | Dfd | g5415.t1 |  |
|  |  |  | Lox5 | g5416.t1 |  |
|  |  |  | Lox4 | g5418.t1 |  |
|  |  |  | Antp | g5417.t1 |  |
|  |  |  | Post2 | g5419.t1 |  |
| *Phoronopsis harmeri* | Pha | Phoronida | lab |  | this study |
|  |  |  | pb |  |  |
|  |  |  | Hox3 |  |  |
|  |  |  | Dfd |  |  |
|  |  |  | Lox5 |  |  |
|  |  |  | Lox4 |  |  |
|  |  |  | Antp |  |  |
|  |  |  | Post2 |  |  |

**Table S2.** Primers used to clone genes from cDNA libraries

| **gene** | **forward** |  | **reverse** |
| --- | --- | --- | --- |
| lab | AGTCAAAACCTGAATCTCTCACACG | GCCCGAACAAGAACACCGAAG |  |
| pb | CATCCAAGCCACACAGTGAAGC | GGTTTGTCCCGAATAGTCTGGTG |  |
| hox3 | GACAACCGCCTCATCCGATAG | CGACACAGTGACTCCTGACACG |  |
| dfd | TGTCGCCTCCCCATCACTCACCAAG | TCAGAGCATCTATTTGAAAACCTGC |  |
| lox5 | GGACTCGCAGCAAACTTATCTTCAC | TTCCATTTCATCCGCCTGTTC |  |
| antp | GCAGAAAAGAAAAGACCAGGACG | AACCCGCACAGTATTATGACCTTG |  |
| lox4 | CGTCTTGCCCTCGTTAGTTCAC | CTCGTTCTCCACATAAGCAAGGAC |  |
| post2 | TGGGCACCTGTGGCTTACTG | TTTTCTTCCTTTTCATCCGTCG |  |

**Alignment of Hox sequences used for phylogenetic analysis (Fig. 2) in FASTA format:**

>Lana_Lox5

EIGYEQKRTRQTYTRYQTLELEKEFHYNRYLTRRRRIEIAHHLGLTERQIKIWFQNRRMKWKKENNIPKLTGP

>Nano_Lox5

DIGYEQKRTRQTYTRYQTLELEKEFHYNRYLTRRRRIEIAHALGLTERQIKIWFQNRRMKWKKENNIAKLTGP

>Pau_Lox5

EIGYEQKRTRQTYTRFQTLELEKEFHYNRYLTRRRRIEIAHALGLTERQIKIWFQNRRMKWKKENNVPKLTGP

>Pha_Lox5

EIGYEQKRTRQTYTRFQTLELEKEFHYNRYLTRRRRIEIAHSLGLTERQIKIWFQNRRMKWKKENNVPKLTGP

>Ttra_Lox5

DIGYEQKRTRQTYTRYQTLELEKEFHFNRYLTRRRRIEIAHALGLTERQIKIWFQNRRMKWKKENNLPKLTGP

>Ctel_Lox5

DFGYEQKRTRQTYTRYQTLELEKEFHYNRYLTRRRRIEIAHALQLTERQIKIWFQNRRMKYKKENNISKLTGP

>Mala_Lox5

EMPIEQKRTRQTYTRYQTLELEKEFHFNKYLTRRRRIEIAHALGLTERQIKIWFQNRRMKWKKENNLQKLTGP

>Ngen_Lox5

EMPIEQKRTRQTYTRYQTLELEKEFHFNKYLTRRRRIEIAHALGLSERQIKIWFQNRRMKWKKENNLQKLTGP

>Esco_Antp

QYGPHRKRGRQTYTRYQTLELEKEFHFNRYLTRRRRIEIAHALCLTERQIKIWFQNRRMKWKKENKAEMPGTE

>Mala_Antp

QFGPDRKRGRQTYTRYQTLELEKEFHFNKYLTRRRRIEIAHALCLTERQIKIWFQNRRMKWKKENKPSEGGTS

>Ngen_Antp

QFGPDRKRGRQTYTRFQTLELEKEFHFNKYLTRRRRIEIAHALCLTERQIKIWFQNRRMKWKKENKQPNGPNC

>Cgig_Lox5

EVTYEQKRTRQTYTRYQTLELEKEFHFNRYLTRRRRIEIAHLLGLTERQIKIWFQNRRMKWKKDNNIPKLTGP

>Esco_Lox5

ETAYEQKRTRQTYTRFQTLELEKEFHFNRYLTRRRRIEIAHSLGLSERQIKIWFQNRRMKWKKENNVSKLTGP

>Gvar_Hox7

DVHFEQKRTRQTYTRYQTLELEKEFHFNRYLTRRRRIEVAHMLGLTERQIKIWFQNRRMKWKKENNVSKLTGP

>Gvar_Lox5

DVHFEQKRTRQTYTRYQTLELEKEFHFNRYLTRRRRIEVAHMLGLTERQIKIWFQNRRMKWKKDNNVSKVTGP

>Ngen_ScrA

---IE-KRTRTSYTRYQTLELEKEFHFNRYLTRRRRIEIAHALNLTERQIKIWFQNRRMKWKKEQKLAHITKS

>Ngen_ScrB

---IE-KRTRTSYTRYQTLELEKEFHFNRYLTRRRRIEIAHALNLTERQIKIWFQNRRMKWKKEQKLAHITKS

>Mala_Scr

---MESKRTRTSYTRYQTLELEKEFHFNRYLTRRRRIEIAHALNLTERQIKIWFQNRRMKWKKEQKLAHITKS

>Lana_Scr

---IESKRTRTSYTRHQTLELEKEFHFNRYLTRRRRIEIAHALNLTERQIKIWFQNRRMKWKKEQKLAHLTKT

>Nano_Scr

----DNKRTRTSYTRHQTLELEKEFHFNRYLTRRRRIEIAHALNLTERQIKIWFQNRRMKWKKEQKLAHLTKT

>Ttra_Scr

--NAESKRTRTSYTRHQTLELEKEFHFNRYLTRRRRIEIAHALNLTERQIKIWFQNRRMKWKKEQKVSHITKN

>Ctel_Scr

---ADNKRTRTSYTRHQTLELEKEFHFNRYLTRRRRIEIAHSLNLTERQIKIWFQNRRMKWKKEHKLAHLAKS

>Ctur_Lox5

--GYEQKRTRQTYTRYQTLELEKEFHYNRYLTRRRRIEIAHTLGLTERQIKIWFQNRRMKWKKENNIAKLTG-

>Fenf_Hox6

---FDRKRGRQTYTRYQTLELEKEFHFNRYLTRRRRIDIAHALCLTERQIKIWFQNRRMKWKKEQKAALGVGM

>Obim_Antp

---PHRKRGRQTYTRYQTLELEKEFHFNRYLTRRRRIEIAHALCLTERQIKIWFQNRRMKWKKENKAEVPVSE

>Fenf_Hox5

DLGIDQKRTRQTYTRHQTLELEKEFHFNRYLTRRRRIEIVHALGLTERQIKIWFQNRRMKWKKENNLKSINDA

>Nano_Dfd

YNGLEPKRSRTAYTRHQILELEKEFHFNRYLTRRRRIEIAHALCLTERQIKIWFQNRRMKWKKEHKLPNTKTR

>Ttra_Dfd

YNGLEPKRSRTAYTRHQILELEKEFHFNRYLTRRRRIEIAHALCLTERQIKIWFQNRRMKWKKEHKLPNTKTR

>Pau_Dfd

YNGLESKRTRTAYTRHQILELEKEFHFNRYLTRRRRIEIAHALCLTERQIKIWFQNRRMKWKKEHKLPNTKTR

>Pha_Dfd

YNGMESKRTRTAYTRHQILELEKEFHFNRYLTRRRRIEIAHALCLTERQIKIWFQNRRMKWKKEHKLPNTKTR

>Lana_Dfd

YNGLEPKRSRTAYTRHQILELEKEFHFNRYLTRRRRIEIAHSLCLTERQIKIWFQNRRMKWKKEHKLPNTKNK

>Mala_Dfd

FNGGENKRTRTAYTRHQILELEKEFHFNRYLTRRRRIEIAHALCLTERQIKIWFQNRRMKWKKEHKLPNTKLR

>Ngen_DfdA

FSSGENKRTRTAYTRHQILELEKEFHFNRYLTRRRRIEIAHALCLTERQIKIWFQNRRMKWKKEHKLPNTKLR

>Ngen_DfdB

FSSGENKRTRTAYTRHQILELEKEFHFNRYLTRRRRIEIAHALCLTERQIKIWFQNRRMKWKKEHKLPNTKLR

>Cgig_Hox4

SLVSESKRNRTAYTRHQILELEKEFHFNRYLTRRRRIEIAHTLCLSERQIKIWFQNRRMKWKKEHKLPNTKTR

>Ctel_Dfd

----DSKRTRTAYTRHQILELEKEFHFNRYLTRRRRIEIAHTLCLSERQIKIWFQNRRMKWKKEHKLPNTKTR

>Fenf_Hox4

NFAGEPKRARTAYTRHQVLELEKEFHFNRYLTRRRRIEIAHALCLTERQIKIWFQNRRMKWKKDHKLPNTKTV

>Lana_Antp

---PDRKRGRQTYSRHQTLELEKEFHFNRYLTRRRRIEIAHALCLTERQIKIWFQNRRMKWKKENKGIELSRD

>Nano_Antp

QFGPDRKRGRQTYSRHQTLELEKEFHFNRYLTRRRRIEIAHALCLTERQIKIWFQNRRMKWKKENRGADLQRD

>Ttra_Antp

G-GPDRKRGRQTYSRHQTLELEKEFHFNRYLTRRRRIEIAHALCLTERQIKIWFQNRRMKWKKENKQEESMKN

>Ctur_Dfda

--GLDPKRARTAYTRHQILELEKEFHFNRYLTRRRRIEIAHTLDLSERQIKIWFQNRRMKWKKEHKLPNTKGK

>Esco_Scr

P-DGESKRSRTSYTRHQTLELEKEFHYNKYLTRRRRIEIAHALNLTERQIKIWFQNRRMKWKKEHKLSHIAKN

>Obim_Scr

P-DGESKRSRTSYTRHQTLELEKEFHYNKYLTRRRRIEIAHALNLTERQIKIWFQNRRMKWKKEHKLSHIAKN

>Gvar_Hox5

GNDADSKRSRTSYTRHQTLELEKEFHYNKYLTRRRRIEIAHALNLTERQIKIWFQNRRMKWKKDHKLSHIAKN

>Ctel_Antp

KAGPERKRGRQTYTRYQTLELEKEFHFNRYLTRRRRIEIAHALCLTERQIKIWFQNRRMKWKKENRQIEVLRQ

>Fenf_Hox8

----PRKRGRQTYTRYQTLELEKEFHFNRYLTRRRRIEMAHALCLTERQIKIWFQNRRMKEKKEKQKIEEMKV

>Mala_Lox4

PNSAQRRRGRQTYSRYQTLELEKEFQFNHYLTRRRRIEIAHSLCLTERQIKIWFQNRRMKLKKERQQIKELND

>Ngen_Lox4

PNSAQRRRGRQTYSRYQTLELEKEFQFNHYLTRRRRIEIAHSLCLTERQIKIWFQNRRMKLKKERQQIKELND

>Ttra_Lox4

PNSAQRRRGRQTYSRYQTLELEKEFQFNHYLTRKRRIEVAHALCLTERQIKIWFQNRRMKLKKERQQIKELND

>Cgig_Lox4

PNSAHRRRGRQTYSRYQTLELEKEFQFNHYLTRKRRIEVAHSLCLTERQIKIWFQNRRMKLKKERQAIKEIND

>Ctel_Lox4

PNSSQRRRGRQTYSRYQTLELEKEFQFNHYLTRKRRIEIAHALCLTERQIKIWFQNRRMKLKKERQQIKDLN-

>Pha_Lox4

PNSAQRRRGRQTYTRYQTLELEKEFQFNHYLTRKRRIEIAHVLCLTERQIKIWFQNRRMKLKKERQQIKEMNE

>Pau_Lox4

PNSAQRRRGRQTYSRYQTLELEKEFQFNNYLTRKRRIEIAHTLRLTERQVKIWFQNRRMKLKKEKQQIKEIN-

>Nano_Lox4

PNSAQRRRGRQTYSRFQTLELEKEFQFNHYLTRKRRIEVAHALCLTERQIKIWFQNRRMKLKKERQQIKEMNE

>Cgig_Lox2

PNSNQRRRGRQTYTRFQTLELEKEFKFNRYLTRRRRIELSHMLCLTERQIKIWFQNRRMKEKKELQAIKELNE

>Ctel_Lox2

PNSNQRRRGRQTYTRYQTLELEKEFKFNRYLTRRRRIELSHMLCLTERQIKIWFQNRRMKEKKEIQAIKELNE

>Obim_Lox2

PNSNQRRRGRQTYTRFQTLELEKEFKFNRYLTRRRRIELSHMLCLTERQIKIWFQNRRMKEKKELQAIKELNE

>Ngen_Lox2

PNSNQRRKGRQTYSRYQTLELEKEFKFNRYLTRRRRVELSALLCLSERQIKIWFQNRRMKEKKEMQAIKELNK

>Gvar_Lox2

QKSNQRRRGRQTYTRFQTLDLEKEFKFNRYLTRRRRIELSHMLCLTERQIKIWFQNRRMKEKKELQAIKELNS

>Gvar_Lox4

PNSAQRRRGRQTYSRYQTLELEKEFQFNHYLTRKRRIEIAHTLCLTERQIKIWFQNRRMKMKKERQAIKDING

>Esco_Lox4

PNSSQRRRGRQTYSRFQTLELEKEFQYNNYLTRKRRIEVAHALNLSERQVKIWFQNRRMKLKKEKQQIREMNG

>Obim_Lox4

PNSSQRRRGRQTYSRFQTLELEKEFQYNNYLTRKRRIEVAHALNLSERQVKIWFQNRRMKLKKEKQQIRELNV

>Gvar_Hox4

--NGEYKRTRTAYTRHQVLELGKEFHFNRYLIRRRRIEITHTLCLSERQIKIWFQNRRMKWKKEHKLPNTKTG

>Pha_Antp

QSAEKKRPGRQTYSRFQTLELEKEFHFNRYLTRRRRVEIAHALGLTERQVKIWFQNRRMKWKKECKALKNMND

>Pau_Antp

--TEKKRVGRQTYTRFQTLELEKEFHFNRYLTRRRRVEIAQVLGLTERQVKIWFQNRRMKWKKEMKALKSLNE

>Ctur_Dfdb

--DGDNKRTRTAYTRQQVLELEKEFHYNRYLTQRRRIEIAHTLTLSERQIKIWFQNRRMKWKKDHKLSSSKGR

>Obim_Lox5

----------------------------------RRIEIAHSLGLSERQIKIWFQNRRMKWKKENNVQKLTGP

>Nano_Hox3

-----SKRARTAYTSAQLVELEKEFHFNRYLCRPRRIEMAALLSLTERQIKIWFQNRRMKFKKEQKQKVLMEK

>Ttra_Hox3

-----SKRARTAYTSAQLVELEKEFHFNRYLCRPRRIEMAALLSLSERQIKIWFQNRRMKFKKEQKQKAILEK

>Lana_Hox3

-----TKRARTAYTSAQLVELEKEFHFNRYLCRPRRIEMAALLNLTERQIKIWFQNRRMKFKKEQKQKVMLEK

>Pau_Hox3

-----AKRARTAYTSAQLVELEKEFHFNRYLCRPRRIEMAALLNLSERQIKIWFQNRRMKFKKEQKQKIAVAK

>Pha_Hox3

-----AKRARTAYTSAQLVELEKEFHFNRYLCRPRRIEMAALLNLSERQIKIWFQNRRMKFKKEQKQKVALEK

>Mala_Hox3

-----PKRSRTAYTSAQLVELEKEFHFNRYLCRPRRIEMAALLNLSERQIKIWFQNRRMKYKKDQKQKNLMEK

>Ngen_Hox3A

-----PKRSRTAYTSAQLVELEKEFHFNRYLCRPRRIEMAALLNLSERQIKIWFQNRRMKYKKDQKQKNLMEK

>Ngen_Hox3B

-----PKRSRTAYTSAQLVELEKEFHFNRYLCRPRRIEMAALLNLSERQIKIWFQNRRMKYKKDQKQKNLMEK

>Ctel_Hox3

-----SKRARTAYTSAQLVELEKEFHFNRYLCRPRRIEMAALLNLTERQIKIWFQNRRMKYKKDQKQKNLMEK

>Cgig_Hox3

--PEPTKRARTAYTSAQLVELEKEFHFNRYLCRPRRIEMAALLSLTERQIKIWFQNRRMKFKKEQRQKPHSEK

>Esco_Hox3

--EQPAKRARTAYTSAQLVELEKEFHFNQYLCRPRRIEMAALLNLSERQIKIWFQNRRMRFKKEKKLKVNMDK

>Gvar_Hox3

--VEPATRARTAYTSAQLVELEKEFHFNRYLCRPRRIEMAALLNLSERQIKIWFQNRRMKFKKDCRLKGGSDK

>Ngen_PbA

--GNNPRRLRTAYTNSQLLELEKEFHFNKYLCRPRRIEIAAALDLTERQVKVWFQNRRMKFKRQTGKGDKGEG

>Ngen_PbB

--GNNPRRLRTAYTNSQLLELEKEFHFNKYLCRPRRIEIAAALDLTERQVKVWFQNRRMKFKRQTGKGDKGDG

>Mala_Pb

--GNNPRRLRTAYTNSQLLELEKEFHFNKYLCRPRRIEIAASLDLTERQVKVWFQNRRMKYKRQSGK----DG

>Lana_Pb

--KQHTRRLRTAYTNTQLLELEKEFHFNKYLCRPRRIEIAASLDLTERQVKVWFQNRRMKFKRQSQLKQQDAP

>Nano_Pb

--SNHARRLRTAYTNTQLLELEKEFHFNKYLCRPRRIEIAASLDLTERQVKVWFQNRRMKFKRQSQLKHGDGS

>Gvar_Hox2

--AGCSRRLRTAYTNTQLLELEKEFHFNKYLCRPRRIEIAASLDLTERQVKVWFQNRRMKYKRQSQS---GRS

>Ttra_Pb

--PNHPRRLRTAYTNTQLLELEKEFHFNKYLCRPRRIEIAASLDLTERQVKVWFQNRRMKFKRQSQNKSDGSG

>Cgig_Hox2

--GGGTRRLRTAYTNTQLLELEKEFHFNKYLCRPRRIEIAASLDLTERQVKVWFQNRRMKYKRQTQSQRQKAE

>Pha_Pb

--CTSQRRLRTAYTNTQLLELEKEFHFNKYLCRPRRIEIAASLDLTERQVKVWFQNRRMKFKRQKQG------

>Ctel_Pb

--SNHPRRLRTAYTNTQLLELEKEFHFNKYLCRPRRIEIAASLDLTERQVKVWFQNRRMKFKRQTGKGSGNSP

>Pau_Pb

--AISQRRLRTAYTNTQLLELEKEFHFNKYLCRPRRIEIAASLDLSERQVKVWFQNRRMKFKRQKQAGHGNRK

>Ctel_Xlox

QTFSENKRTRTAYTRAQLLELEKEFHFNRYITRPRRVELAAHLNLTEQHIKIWFQNRRMKWKKDVDKKRPQQS

>Fenf_MedPost

---TPHRERRQTYTRHQTAELEREYVTNRYLTRRRRIEISQSLHLSERQIKIWFQNRRMKEKREKDHVTSPRK

>Mala_Lab

--AGQPNTGRTNFTNKQLTELEKEFHFNKYLTRARRIEIAAALGLNETQVKIWFQNRRMKQKKRMKEGLVQNN

>Ngen_LabA

--AGQPNTGRTNFTNKQLTELEKEFHFNKYLTRARRIEIAAALGLNETQVKIWFQNRRMKQKKRMKEGLVQNN

>Ngen_LabB

--AGQPNTGRTNFTNKQLTELEKEFHFNKYLTRARRIEIAAALGLNETQVKIWFQNRRMKQKKRMKEGLVQNN

>Lana_Lab

--GNIPNMGRTNFSNKQLTELEKEFHFNKYLTRARRIEIAAALGLNETQVKIWFQNRRMKQKKRMKESQTLP-

>Nano_lab

--ANAPNMGRTNFSNKQLTELEKEFHFNKYLTRARRIEIAAALGLNETQVKIWFQNRRMKQKKRLKEGGSMT-

>Ttra_Lab

--NQAPNMGRTNFSNKQLTELEKEFHFNKYLTRARRIEIAAALGLNETQVKIWFQNRRMKQKKRMKESPLSAH

>Pau_Lab

--SAASNMGRTNFTNKQLTELEKEFHFNKYLTRARRIEIAATLGLNETQVKIWFQNRRMKQKKRMKECRAQMK

>Pha_Lab

--TTATNMGRTNFTNKQLTELEKEFHFNKYLTRARRIEIAATLGLNETQVKIWFQNRRMKQKKRMKECRAQMK

>Ctel_Lab

--AGQPNMGRTNFTNKQLTELEKEFHFNKYLTRARRIEIAASLGLNETQVKIWFQNRRMKQKKRLKENTSTTP

>Fenf_Hox1

--GGINNTGRTNFTTKQLTELEKEFHFNKYLTRARRIEIAGALQLNETQVKIWFQNRRMKQKKRMKEGLIPPD

>Gvar_Hox1

--GGINSTGRTNFWNKQAFEFEKEFHFNKYFTRARRIEIAAALGLNETQVKIWFQNSRMKQKKRMSEIQFEKG

>Cgig_Hox1

--TPQPNMGRTNFTNKQLTELEKEFHFNKYLTRARRIEIAAALGLNETQVKIWFQNRRMKQKKRLREAQFDQN

>Obim_Hox1

--VGGGGTGRTNFTNKQLTELEKEFHFNKYLTRARRIEIAAALGLNETQVKIWFQNRRMKQKKRLKEAQGTTG

>Esco_Lab

--GGGNSTGRTNFTNKQLTELEKEFHFNKYLTRARRIEIAAA-------------------------------

>Pha_Xlox

--------------------LEKEFHFNKYISRPRRIELAALLNLTERHIKIWFQNRRMKWKKDEAKRRPRPL

>Ctel_Gsx

ESSDAVKRMRTAFSSTQLLELEREFASNMYLSRLRRIEIATYLSLSEKQVKIWFQNRRVKFKKEGAAHGSRDH

>Pha_Gsx

DDLDSSKRIRTAFTSTQLLELEREFAANMYLSRLRRIEIATYLNLSEKQVKIWFQNRRVKYKKEGG--GSRDR

>Ctel_Eve

------RRYRTAFTREQLGRLEREFLKENYVSRPRRCELAASLNLPESTIKVWFQNRRMKDKRQRLAMA----

>Esco_Post1

PSAIALRKRRRPYSKYQIAELEREYALSTYISKSRRWELSQLLNLSERQIKIWFQNRRIKAKKLQKRDETLKT

>Obim_Post1

PSAIALRKRRRPYSKYQIAELEREYAISTYISKSRRWELSQLLNLSERQIKIWFQNRRIKAKKLQKRDETLKG

>Gvar_Post1

PTTVTLRKRRRPYSKFQIAELEREYN-GSYVSESRRWELSQLINLSERQIKIWFQNRRIKAKKIIKRDDISPQ

>Lana_Post1

PAVIHMRKKRKPYSKYQIAELEREYVSNTYISKPKRWELSQRLQLSERQVKIWFQNRRMKEKKVKGGKQT---

>Ttra_Post1

PTVIHMRKRRKPYSKQQINELEREYVKTTYISKPKRWELAQRLNLSERQVKIWFQNRRMKEKKMRGSRRX---

>Ctur_Pb

------------------------FHFNKYLCRPRRIEIAASLDLTERQVKV---------------------

>Ctur_Hox3

------------------------FHFNRYLCRPRRIEMAALLSLTERQIKI---------------------

>Ngen_Post2A

---PRTRKKRKPYTRYQTMVLENEFMTNSYITRQKRWEISCKLHLSERQVKVWFQNRRMKRKKLNARTK-IKS

>Ngen_Post2B

---PRTRKKRKPYTRYQTMVLENEFMTNSYITRQKRWEISCKLHLSERQVKVWFQNRRMKRKKLNARTK-IKS

>Mala_Post2

---PRTRKKRKPYTRYQTMVLENEFMTNSYITRQKRWEISCKLHLTERQVKVWFQNRRMKRKKLNSRAK-VKS

>Ctel_Post2

---PKQRKKRKPYTRYQTMVLENEFINNSYITRQKRWEISCKLHLSERQVKVWFQNRRMKRKKLNERAK---S

>Esco_Post2

---TKGRKKRKPYTRYQTMVLENEFLNSSYITRQKRWEISCKLQLTERQVKVWFQNRRMKRKKLNERAK-ARL

>Obim_Post2

---TKGRKKRKPYTRYQTMVLENEFLNSSYITRQKRWEISCKLQLTERQVKVWFQNRRMKRKKLNERAK-ARL

>Pau_Post2

-----RGKKRKPYTRYQNMVLENEFLSSSYITRQKRWEISCKLHLTERQVKVWFQNRRMKRKKINERAK-ALF

>Pha_Post2

---IRTRKKRKPYTRYQNMVLENEFLGSSYITRQKRWEISCKLHLTERQVKVWFQNRRMKRKKINERAK-ALF

>Lana_Post2

---AKQRKKRKPYTRYQTMVLENEFLNNAYITRQKRWEISCKLHLSERQVKVWFQNRRMKRKKLNERAK-ALF

>Nano_Post2

---IRTRKKRKPYTRYQTMVLENEFLNSAYITRQKRWEISCKLHLSERQVKVWFQNRRMKRKKLNERAK-ALF

>Ttra_Post2

---FRSRKKRKPYTRYQNMLLENEFIANSYITRQKRWEISCKLQLTERQVKVWFQNRRMKRKKLTDRAK-SLF

>Ctur_Post2

---SSSRKKRKPYTRYQTMVLETEFINNSYITRQKRWEISCRLRLTERQVKVWFQNRRMKRKKLNDRAKNAQL

>Fenf_Hox3

-------------------------HFNRYLCRPRRIEMAALLNLSERQIKI---------------------

>Fenf_PostA

------RKKRKPYTKHQTFILEQEYLMSTYITRQRRLELARNLSLTERQVKIWFQNRRMKTKKLRERNKVSVG

>Fenf_PostB

TSGGKCRTKRKPYEKWVTYLLEEEYLSNTYITKQKRYELSYRTSLTERQVKIWFQNRRMKSKKLRERSTSGAT

>Ctel_Cdx

GKTRTKDKYRIVYSEYQKVELEKEYLYSKYITIQRKAELSRSIGLSERQVKIWFQNRRAKERKQKRKMEEALS

>Ctel_Post1

---VNPKKKRKPYSKPQVSALENEYSTSTYITKARRKEVARELDLTERQIKIWYQNRRIKEKKIATKRAKVQS
